# DNA barcoding reveals an increased diversity within the genus *Culex* (Diptera: Culicidae) in Ireland

**DOI:** 10.1101/2023.12.20.572570

**Authors:** Thomas G. Curran, Aidan O’Hanlon, Finán Gallagher, Rasmus S. Pedersen, Annetta Zintl, Elsie Isiye, Angela Valcarcel Olmeda, David O’Neill, Allan D. McDevitt, Catherine O’Reilly, Denise B. O’Meara

## Abstract

Arthropod populations are changing globally due to the combined impacts of climate change, land-use transformation, globalisation and trade. Changes in arthropod populations can potentially result in the decline and extinction of some native and/or endemic species, while other species that potentially pose threats may increase. Such dynamics lead to weakened ecosystem resilience, which can facilitate the establishment and spread of species with vector capabilities, such as mosquitoes. While many of the mosquito species present in Ireland possess vector capabilities, populations have not been nationally surveyed since 1991. At that time, species identification relied on morphological examination, a method that was even then recognised as inadequate for differentiating certain species within complexes. In this study, we present the first DNA barcoding investigation of Irish mosquito populations and reveal species and hidden diversity not previously recognised in Ireland. These include two potential new species/ecotypes, *Culex torrentium* and *Culex pipiens* form *pipiens*, which were recorded for the first time in Ireland. We demonstrate that barcoding with Cytochrome C Oxidase Subunit 1 (COI) based primers is suitable for the majority of species-level identifications. However, additional primer sets targeting the Internal Transcribed Spacer regions 1 and 2 (ITS1 and ITS2) were necessary to differentiate species within the *Culex pipiens* complex. The multi-locus approach employed in this study can enhance national surveillance efforts, especially in monitoring mosquitoes that may transmit vector-borne diseases, and in recognising increased species diversity from a biodiversity perspective.

## Introduction

In Britain, 36 species of mosquitoes have been described (Supplementary Table 1) (Cranston et al. 1987; Kampen et al. 2015; Vaux and Medlock 2015), and ongoing monitoring of mosquitoes has resulted in the continuous expansion of the British mosquito checklist. An entomology service for the identification of mosquitoes submitted by environmental health officers has also been established (Kampen et al. 2015; Vaux and Medlock 2015). In Ireland, the only active surveillance for mosquitoes is conducted by environmental health officers from the Health Service Executive (HSE). This surveillance focuses on potential points of entry for exotic species, such as airports and seaports, and to our knowledge the data are unpublished. The last published Irish mosquito survey was conducted by Ashe et al. (1991), who documented 18 species, including the *Anopheles maculipennis* complex (Supplementary Table 1), a complex which is thought to be comprised of 11 species; *An. artemievi*, *An. atroparvus*, *An. beklemishevi*, *An. daciae*, *An. labranchiae*, *An. maculipennis*, *An. martinius*, *An. melanoon*, *An. messeae*, *An. persiensis,* and *An. sacharovi* (Fantini 1994; White 1978; Linton et al. 2007; Kampen et al. 2016; Kavran et al. 2018). At the time of the survey it was suggested that *An. atroparvus* and *An. messeae* may also occur in Ireland, but it was not possible to confirm these identifications based on the morphology of the two species (Ashe et al. 1991) implying that even then, the number of mosquito species in Ireland was thought to be underestimated. Highlighting the need for the development of multi-faceted approaches to enable exact mosquito species identification, which can complement traditional morphological techniques.

Recently, Curran et al. (2022) used DNA metabarcoding of an insectivorous predator, the lesser horseshoe bat (*Rhinolophus hipposideros*), to investigate its diet across the west of Ireland. In doing so, four mosquito species were potentially detected: *Culiseta annulata*, *Cs. morsitans*, *Culex pipiens*, and *Cx. quinquefasciatus*. The latter of these species was not previously reported in Ireland and suggests that Irish mosquitoes, particularly those within the genus *Culex*, may be more diverse than previously thought. However, the sequence regions utilised in Curran et al. (2022) were notably short, spanning separate 133 bp and 157 bp fragments of the Cytochrome C Oxidase Subunit 1 gene (Browett et al. 2021). Curran et al. (2022) noted that further studies using longer sequence regions are necessary for accurate species identification.

The *Culex pipiens* complex remains enigmatic in both Ireland and Britain. In Ireland, only one species of *Culex* has been documented, *Cx. pipiens* (Ashe et al. 1991), while four *Culex* species are recognised in Britain (*Cx. modestus*, *Cx. pipiens*, *Cx. territans*, and *Cx. torrentium*). Members of the *Cx. pipiens* complex are difficult to distinguish morphologically, but differ in terms of behaviour, physiology, and host preference (Zittra et al. 2016). Depending on the source, the *Cx. pipiens* complex comprises a variable number of species; *Cx pipiens* and its associated ecotypes (*Cx. pipiens* form *pipiens* and *Cx. pipiens* form *molestus*), *Cx. quinquefasciatus*, *Cx. torrentium*, *Cx. pipiens pallens*, *Cx. vagans*, *Cx. australicus*, and *Cx. globocoxitus* (Vinogradova 2000; Zittra et al. 2016; Aardema et al. 2020). An ecotype is described as intraspecific groups of species with distinguishable and heritable differences allowing adaptation and subsequent prevalence to local environmental conditions (Lowry 2012; Wadgymar et al. 2022; Small et al. 2023). The two ecotypes representing *Cx. pipiens* (i.e. *Cx. pipiens* form *pipiens* and *Cx. pipiens* form *molestus*) can be differentiated using molecular techniques and in Northern Europe microsatellite studies have reported unique fingerprints for these two ecotypes and it was implied that this is indicative of separate species status. However, there is still much debate about the exact taxonomic status for these ecotypes and to date they are regarded as subspecific forms of *Cx. pipiens* (Wilkerson et al. 2021). Based on this, these ecotypes will be referred to as *Cx. pipiens* form *pipiens* and *Cx. pipiens* form *molestus* throughout.

A range of arthropod-borne diseases (arboviruses) are associated with the *Cx. pipiens* complex (Becker et al. 2010; Dörge et al. 2020; Hesson et al. 2014; Werblow et al. 2013). Of these, West Nile Virus (WNV) is most notably linked to *Cx. pipiens*. This arbovirus has caused infections and epidemics throughout Europe (Hayes 2001; Hubálek 2008; Lundström 1999). In Ireland, one case of WNV was diagnosed in 2012 in a person who had acquired it in the United States. However, it is important to stress that up to 80% of cases may be asymptomatic, highlighting the need to understand resident mosquito vector capabilities (HPSC – Ireland’s Health Protection Surveillance Centre https://www.hpsc.ie/a-z/vectorborne/westnilevirus/epidemiologicaldata/).

This paper describes the use of DNA barcoding to identify field-collected mosquitoes. The aim of the study was to develop and optimise methods for the DNA-based identification of mosquito species occurring in Ireland, using larval and adult specimens. We employed DNA barcoding with primer sets targeting the Cytochrome C Oxidase Subunit 1 (COI) gene and Internal Transcribed Spacer Regions 1 and 2 (ITS1/2) to determine the most suitable genetic locus for species-level identification across various species, and phylogenetic analysis was conducted to explore mosquito diversity in Ireland.

## Results

### DNA Extractions

A minimally destructive approach was used to acquire DNA from adults using just two legs to ensure an almost intact specimen was available for any morphological assessments that may be required, but this resulted in a much lower concentration of DNA when compared to using a whole larva (Supplementary Table 2). Average larval DNA concentration was 93.99 (± 74.66) ng/µL, while the average DNA concentration for adult samples was 8.8 (± 4.41) ng/µL. Although larval samples, on average, yielded a higher concentration of DNA, high levels of variations were observed in the larval samples, and while the NMI samples yielded the lowest concentration of DNA in comparison to other larval samples, the results were more consistent.

### DNA barcoding with COI and ITS1

All specimens were successfully amplified via PCR for both the COI and ITS1 loci. Out of the 84 samples, 82 provided successful sequences via COI barcoding and 81 via ITS1 barcoding. Supplementary Table 3 lists the species identified from each sample, along with the percentage identity, percentage query cover, and the accession number of the top BLAST identification from GenBank. Three and four genera of mosquitoes were identified using either the COI or the ITS1 genetic region, respectively. Both COI and ITS1 detected *Cx. pipiens*, *Cx. torrentium*, and *Cx. quinquefasciatus*. Additionally, ITS1 facilitated the identification of ecotypes/hybridised species within *Cx. pipiens*, namely *Cx. pipiens* form *pipiens* (N = 4) (percentage identification: 98.75% - 99.87% ± 0.46%; query cover: 99% - 100% ± 0.43%) and *Cx. pipiens pallens* (N = 2) (percentage identification: 94.44% - 95.30% ± 0.43%; query cover: 99% - 100% ± 0.5%) (Table 1). Other species identified included *Culiseta annulata* (DA = 3, DL =3) and *Anopheles claviger* (NI = 14). These identifications were based solely on COI barcoding, as ITS1 provided contradictory identifications unsupported by low percentage query covers. Where COI identified samples as *Cs. annulata*, ITS1 suggested either *Cs. novaezealandiae* (query cover: 32% – 33%), *Aedes mcintoshi* (query cover: 25% - 34%), or *Ae. metallicus* (query cover: 33%). Moreover, samples identified as *An. claviger* using the COI gene were identified as *An. algeriensis* (query cover: 33% - 36%) when using ITS1.

**Table 1:**
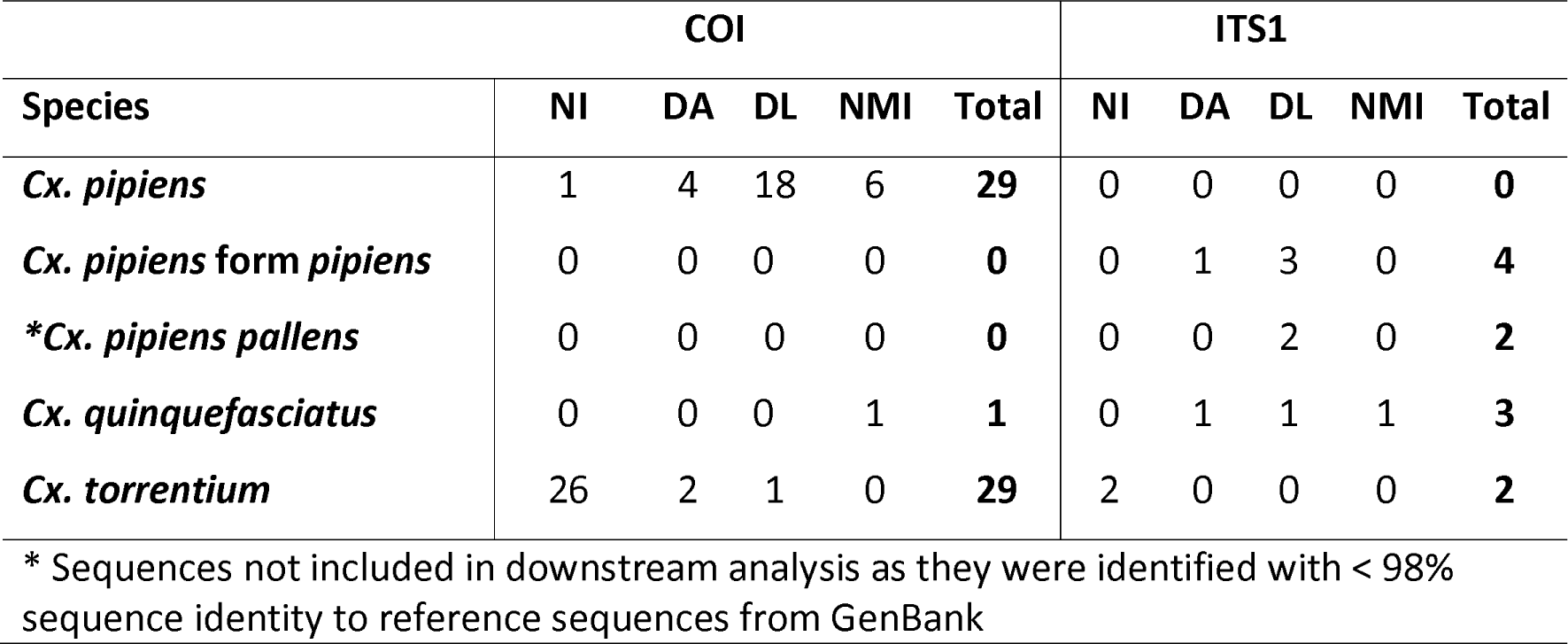
Number of specimens identified within the Genus Culex via COI and ITS1 barcoding with a minimum of 98% sequence identity to a GenBank reference sequence (NI = samples from Warrenpoint, Co. Down; DA = adult samples from Dundalk, Co. Louth; DL = larval samples from Dundalk, Co. Louth; NMI = larval samples sent from the National Museum of Ireland).

### *Culex* Species Identifications based on Phylogenetic Analysis

Fig 1 shows the phylogenetic analysis of specimens in the *Culex* genus based on the COI (A) and ITS1 (B) genetic regions. Firstly, in the COI phylogenetic tree in Fig 1A, samples NI1, NI3, and NI12 from Warrenpoint, Northern Ireland, aligned with *Cx. torrentium*, confirming the DNA BLAST results shown in Supplementary Table 3. A Northern Ireland sample (NI7) and two from Dundalk (DL6 and DL9) appear within a mixed group of sequences, which includes *Cx. quinquefasciatus*, *Cx. pipiens*, *Cx. pipiens* form *pipiens*, *Cx. pipiens* form *molestus*, and *Cx. pipiens pallens*. Meanwhile, two larval samples collected from Irish airports and seaports, and previously identified morphologically as *Cx. territans* at the National Museum of Ireland (NMI) – namely, NMI1 and NMI3 – were genetically identified as *Cx. pipiens*.

**Figure 1:**
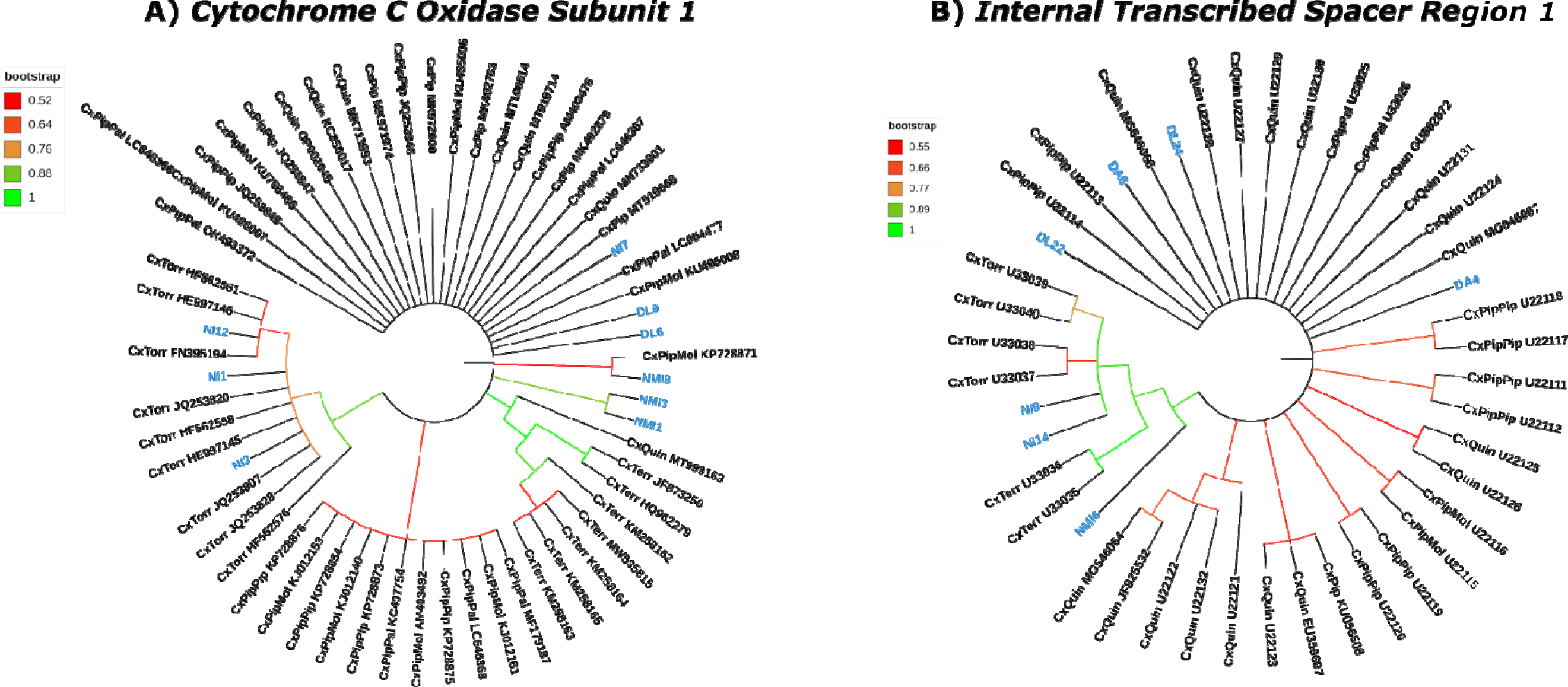
Maximum-Likelihood Phylogenetic Tree (1000 iterations) using the (A) Cytochrome C Oxidase Subunit 1 and (B) Internal Transcribed Spacer Region 1 of samples from Northern Ireland (NI), Dundalk (DA = Adult specimen; DL = Larval specimen) and samples collected at airports and seaports (NMI) molecularly identified as Culex species (highlighted in blue) compared to other Culex species available from GenBank. [CxTorr = Cx torrentium; CxPip = Cx. pipiens; CxQuin = Cx. quinquefasciatus; CxPipPip = Cx. pipiens form pipiens; CxPipPal = Cx. pipiens pallens; CxPipMol = Cx. pipiens form molestus; CxTerr = Cx. territans].

The ITS1 phylogenetic tree in Fig 1B, demonstrates a similar trend. Samples NI9 and NI14 from Warrenpoint, Northern Ireland, aligned with *Cx. torrentium*, again confirming the DNA BLAST results in Supplementary Table 3. Four samples from Dundalk (DA4, DA6, DL22, and DL24) appear within an undifferentiated group of sequences, including *Cx. quinquefasciatus*, *Cx. pipiens pallens*, and *Cx. pipiens* form *pipiens*.

### Culex pipiens complex confirmation

COI and ITS1 barcoding suggested the presence of both *Cx. pipiens* form *pipiens* and *Cx. quinquefasciatus*, but neither primer set provided definitive confirmation (Supplementary Table 3). PCR amplification using the ITS1/2 primers resulted in amplicons that matched *Cx. quinquefasciatus* most closely in two adult specimens (DA4 and DA6) while four larval samples matched *Cx. pipiens* form *pipiens* (Table 2). Nevertheless, the identification of *Cx. quinquefasciatus* was deemed tentative by the authors, given its known distribution in tropical/sub-tropical regions.

**Table 2:**
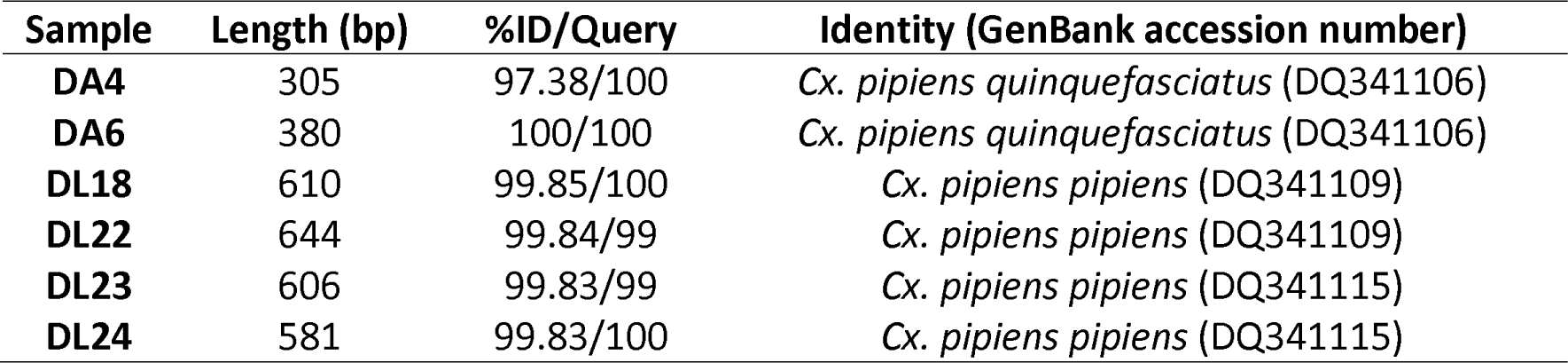
Sanger sequencing results using Culex pipiens complex-specific ITS1/2 primers by Crabtree et al. (1995) of mosquito adults (DA) and larvae (DL) that have been identified as either Culex quinquefasciatus or Culex pipiens form pipiens via ITS1 gene sequencing.

| Sample | Length (bp) | %ID/Query | Identity (GenBank accession number) |
| --- | --- | --- | --- |
| DA4 | 305 | 97.38/100 | *Cx. pipiens quinquefasciatus* (DQ341106) |
| DA6 | 380 | 100/100 | *Cx. pipiens quinquefasciatus* (DQ341106) |
| DL18 | 610 | 99.85/100 | *Cx. pipiens pipiens* (DQ341109) |
| DL22 | 644 | 99.84/99 | *Cx. pipiens pipiens* (DQ341109) |
| DL23 | 606 | 99.83/99 | *Cx. pipiens pipiens* (DQ341115) |
| DL24 | 581 | 99.83/100 | *Cx. pipiens pipiens* (DQ341115) |

In Fig 2, DL23 and *Cx. pipiens* form *pipiens* (accession number: U22112) are shown to group together. DL24 is grouped with *Cx. pipiens* form *pipiens* (accession numbers: DQ341115 and U22111) and *Cx. pipiens* form *molestus* (accession number: U22115). Sample DA6 groups with *Cx. quinquefasciatus* (DQ341106) with a high bootstrap value (bootstrap value = 0.703). Given the branching pattern in Figure 3, sample DA4 appears to have genetic similarities with *Cx. quinquefasciatus*, referencing accession number DQ341106, despite a percentage identity below the 98% criterion.

**Figure 2:**
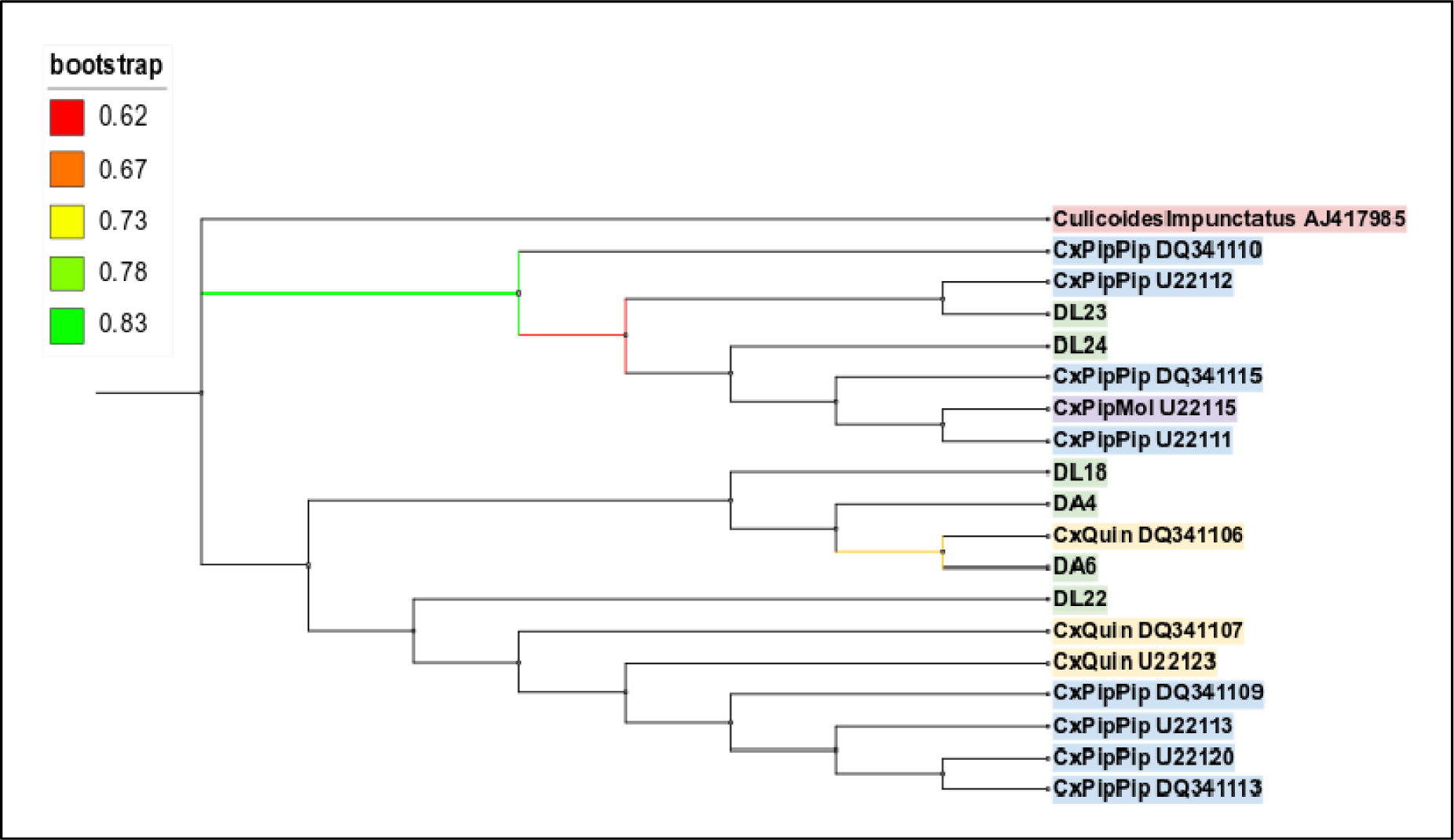
Maximum Likelihood phylogenetic tree (1000 iterations) of specimens identified as Culex quinquefasciatus or Culex pipiens form pipiens (green) compared to sequences from GenBank belonging to Culex quinquefasciatus (yellow), Culex pipiens form pipiens (blue), Culex pipiens form molestus (purple), and an outgroup species Culicoides impunctatus (red) [DL – Dundalk Larva; DA – Dundalk Adult] using a 289 bp alignment of ITS1/2.

**Figure 3:**
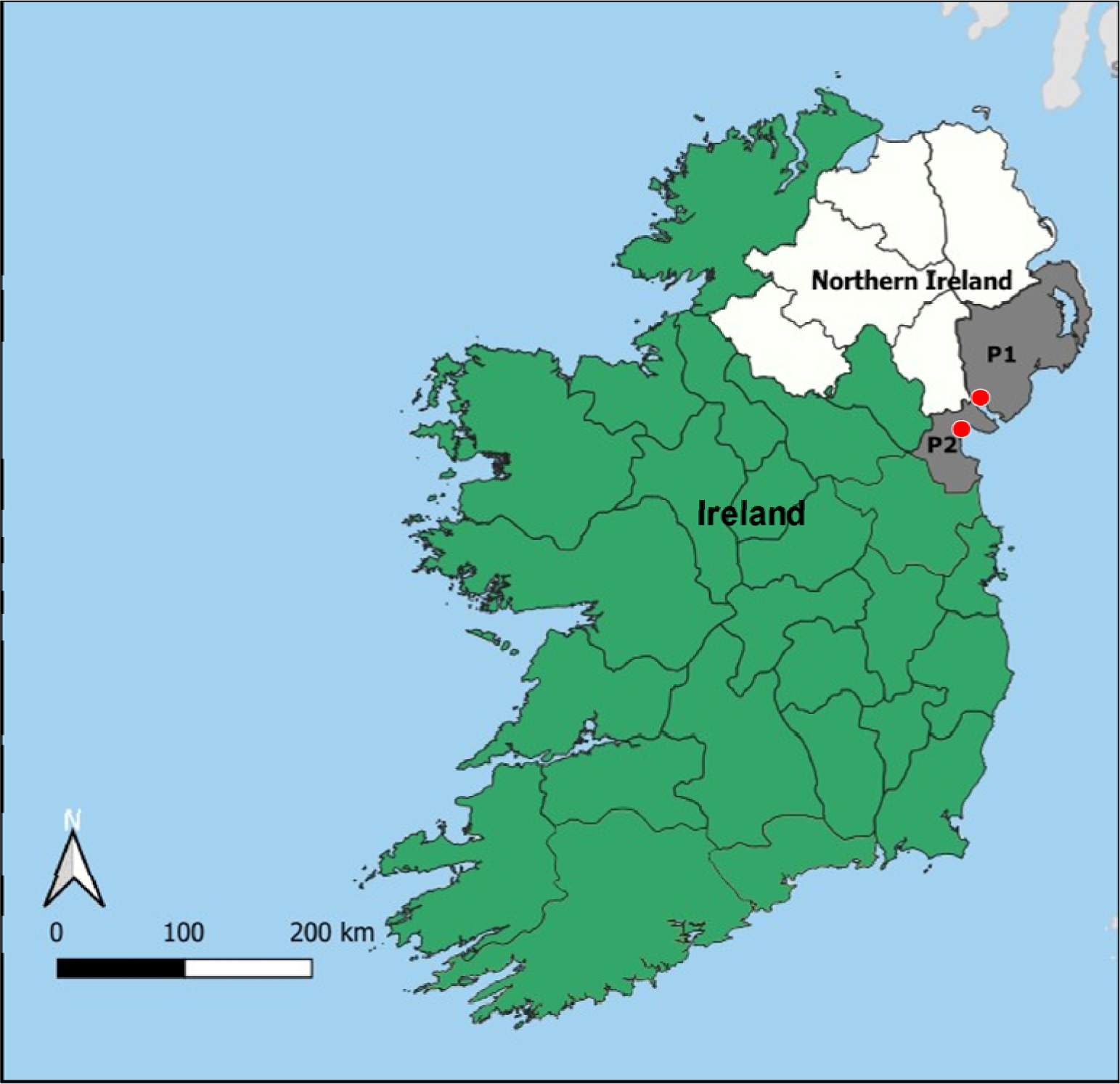
Mosquito sampling. P1: Warrenpoint, Co. Down, Northern Ireland; P2: Dundalk, Co. Louth, Republic of Ireland.

## Discussion

The only mosquito species list currently available in Ireland dates back to the early 1990s and is based only on morphological data (Ashe et al. 1991). This is the first Irish study to employ molecular identification methods to mosquito surveillance and confirmed the presence of *Anopheles claviger*, *Culiseta annulata*, and *Culex pipiens* via COI and ITS DNA barcoding. In addition, three potential new species/ecotypes were recorded for the first time in Ireland, *Cx. torrentium*, *Cx. pipiens* form *pipiens*, and *Cx. quinquefasciatus*, the latter of which is questionable, as discussed below, and requires further investigation.

### Collection of mosquito specimens

The passive collection approach applied in this study facilitated the direct sampling of mosquito adults and larvae and was time- and resource-efficient. The majority of samples collected and analysed in this study were mosquito larvae (n = 75) and it has been suggested that controlling mosquitoes at the larval stages is essential in the prevention of a mosquito-borne disease outbreak (Becker et al. 2010), and here we demonstrate the ease in which mosquito larvae can be collected through the involvement of public participation in mosquito surveys. Furthermore, while it may prove difficult to morphologically identify mosquito larvae to species-level, they are easily distinguishable from other dipteran families in the field, so fresh specimens can be collected from water sources with minimal effort. This involvement, in addition to active surveillance methods, may prove highly valuable in the detection of exotic species in Ireland.

Depending on the sample type, different DNA extractions methods were employed. Larvae typically yielded higher, but more variable, DNA concentrations than adults (for which only two legs were used, preserving the remainder of the specimen for future analysis), and the larval samples supplied by the National Museum of Ireland (NMI) yielded notably lower concentrations of DNA in comparison to those collected and preserved quickly in the field. These NMI samples, collected by environmental health officers from the Health Service Executive (HSE) at potential invasive species entry points, were examined morphologically, and may not have been stored in conditions suitable for downstream DNA analysis. For optimal molecular identification in future studies, we recommend focusing on larval samples, where possible, and extracting DNA promptly to help conserve the quantity and quality of the DNA sample.

### Use of the COI and ITS1 loci for species identification

Both COI and ITS loci have been employed in previous barcoding efforts for mosquito identification (Miller et al. 1996; Rosera-García et al. 2017; Ruiz-Arrondo et al. 2019; Chaiphongpachara et al. 2022). Combining genetic regions from both mitochondrial (COI) and nuclear (ITS) regions is crucial for building comprehensive databases representing mosquito sequences and potential haplotypes. In this study, analysing these regions individually and in parallel has offered significant insights into Irish mosquito diversity. Such depth of understanding was previously unattainable due to the challenges posed by morphological similarities among species. Barcoding of the COI gene allowed for the identification of five species: *An. claviger* (n = 14), *Cs. annulata* (n = 6), *Cx. pipiens* (n = 30), *Cx. quinquefasciatus* (n = 1), and *Cx. torrentium* (n = 29). All specimens identified as *Cx. torrentium* showed a minimum of 98% sequence identity compared to their corresponding GenBank reference sequences. Moreover, all but one of the specimens identified as *Cx. pipiens* satisfied the minimum 98% sequence identity criterion. Intriguingly, one sample (NMI4) matched with *Cx. quinquefasciatus*, exhibiting 100% sequence identity and query cover relative to a specified GenBank sequence (accession number: MK370091). However, a DNA blast of this sequence revealed over 99% similarity to *Cx. pipiens* casting significant doubt regarding the reliability of the identification of that sample.

ITS1 barcoding identified six potential mosquito species/ecotypes, including notable ones like *Cx. torrentium* and *Cx. pipiens* form *pipiens*. The identification of *Cx. torrentium* was reaffirmed here from the same 29 samples identified as that species using COI barcoding, supporting its categorisation within the sample set. Diverging from COI findings, ITS1 identified *Cx. pipiens* in merely two samples, both displaying sequence identities below 98%. In contrast, *Cx. pipiens* form *pipiens* was detected in 22 samples, with five achieving the 98% identity threshold. A further five specimens were identified as *Cx. quinquefasciatus* via ITS1 barcoding, with three surpassing the 98% sequence threshold. A hybrid species resulting from interbreeding of *Cx. pipiens* and *Cx. quinquefasciatus*, known as *Cx. pipiens pallens* (Harbach 2012), was identified in two samples; however, in both cases this identification was dismissed due to insufficient identities (both below the 98% threshold). These tentative *Cx. pipiens pallens* identifications matched accession numbers U33025 and U33026 from GenBank which correspond to a study by Miller et al. (1996) who aimed to resolve the phylogeny of *Culex* species using the two ITS regions (i.e. ITS1 and ITS2). However, a BLAST search found that these sequences were also related to other sequences of *Cx. pipiens* form *pipiens*, *Cx. pipiens* form *molestus*, and *Cx. quinquefasciatus*. In their study, Miller et al. (1996) constructed various phylogenetic trees and *Cx. pipiens pallens* can be seen appearing between ITS sequences of *Cx. pipiens* and *Cx. quinquefasciatus*, demonstrating the lack of genetic resolution to differentiate these species and their hybridised form based on phylogenetic analysis. Additionally, Miller et al. (1996) observed intermediates of *Cx. pipiens/quinquefasciatus* forming a group with *Cx. pipiens pallens* that occurs within the clade of *Cx. pipiens* and stated that this supports evidence that *Cx. pipiens pallens* is a hybridised population of *Cx. pipiens* and *Cx. quinquefasciatus*, rather than the previously proposed subspecies classification. Although we have reservations about our identification of *Cx. quinquefasciatus* and *Cx. pipiens pallens*, our work echoes the complex relationships of species within the *Cx. pipiens* complex.

Regarding the non-*Culex* samples, a disagreement for species-level identifications were observed between COI and ITS1 barcoding (Supplementary Table 3). Samples identified as *An. claviger* using COI were identified as *An. algeriensis* via ITS1 barcoding, both of which are species that have already been documented in Ireland (Ashe et al. 1991). The COI sequences for *An. claviger*, however, demonstrated higher percentage identities and query coverage against GenBank references in comparison to the ITS1 sequence of *An. algeriensis*. In addition to this, while both species were reported in the previous Irish mosquito survey *An. claviger* was one of the more common species documented (Ashe et al. 1991). Thus, the identification of *An. claviger* was preferred over *An. algeriensis*. COI gene sequencing identified *Cs. annulata* from several samples, however this could not be confirmed based on ITS1 sequencing due to the lack of suitable reference data on GenBank, with no ITS1 and only one ITS2 sequence for *Cs. annulata* available. ITS1 barcoding suggested *Ae. mcintoshi*, *Ae. metallicus*, and *Cs. novaezealandiae* in lieu of *Cs. annulata* at very low percentage identities and query covers. Nevertheless, the COI identification of *Cs. annulata* was preferred over ITS1 suggestions due to the species ubiquitous nature in Ireland, as recorded by Ashe et al. (1991), and its detection within the diet of lesser horseshoe bat populations by Curran et al. (2022).

### Identification of new and potentially occurring species in Ireland

The identification of *Cx. torrentium* in Ireland is an expected addition to the nation’s checklist, given its morphological similarities to members of the *Cx. pipiens* complex and its widespread presence across many parts of Europe alongside *Cx. pipiens* (Vinogradova 2000). Intriguingly, this species was not recorded in Britain until 1951 due to the high levels of morphological crossover hindering species identification (Service 1968). Consequently, it has been suggested that due to a conflation with *Cx. torrentium* that *Cx. pipiens* populations might have been overestimated in the past (Hesson et al. 2014). Similarly, Ashe et al. (1991) may have missed the presence of *Cx. torrentium*, given the inherent challenges in distinguishing members of this complex based solely on morphology, but the authors did note that this and seven other mosquito species are expected to occur in Ireland. Although no known vector-related risks are associated with this species in Ireland, laboratory-based studies in Germany have identified it as a competent vector for WNV. In fact, this species was shown to be a more efficient/suitable vector for WNV than both *Cx. pipiens* form *pipiens*, also identified in this study, and *Cx. pipiens* form *molestus* (Jansen et al. 2019), leading the authors to recommend that this species is closely monitored across Central and Northern Europe.

The unexpected identification of *Cx. quinquefasciatus* in Ireland seems questionable. The species, predominantly found in tropical and subtropical regions, has not been reported in Europe, and Ireland lies well outside its current distribution range (Alaniz et al. 2018). This is also reflected in the study locations for the accession numbers which our putative *Cx. quinquefasciatus* COI and ITS1 sequences matched, i.e. Bangladesh, New Zealand, Australia, and Papa New Guinea. It is important to stress that our study’s findings hinge on accurately identified sequences from GenBank and potential mislabelling from previous submissions can lead to further misidentifications. For example, a DNA blast of the COI sequence of the apparent *Cx. quinquefasciatus* sequence from our study aligned more closely with *Cx. pipiens* and *Cx. pipiens* form *pipiens*. While it was not possible to definitively pinpoint *Cx. quinquefasciatus* samples among the other *Culex* species based on ITS1 sequences, the ITS1/2 primers developed by Crabtree et al. (1995) matched a *Cx. quinquefasciatus* sample (i.e. DA6) with 100% identity and query cover across 380bp. However, another DNA blast of the accession number related to this identification (i.e. DQ341106) indicated a close sequence similarity to *Cx. pipiens* in that case too, raising more doubts about the authenticity of our identification. Another difficulty associated with definitive identification of the species is that there are few morphological characteristics that can distinguish *Cx. quinquefasciatus* from other *Culex* species (Harbach 2012). This is further compounded by the presence of hybrids including the hybrid potentially identified in this study, *Cx. pipiens pallens*.

On the other hand, it is interesting to note that models by Samy et al. (2016) and Alaniz et al. (2018) have identified temperate regions, including Ireland, as potentially suitable for *Cx. quinquefasciatus*; findings that are supported by its detection in Turkey, which has European territory (Gunay et al. 2015; Hernández-Triana et al. 2022). In the USA, where *Cx. quinquefasciatus* is an important vector for WNV it is thought to amplify the virus among avian hosts due to its ornithophilic feeding behaviour, and serves as a bridge vector to humans (Adams et al. 2021). In the last 15 years, it has also expanded its distribution in New Zealand. This spread southwards is thought to be facilitated by changes in land use and the environment, and its increasing presence seems to coincide with a decline in a native *Culex* species to New Zealand, *Cx. pervigilans* (Kasper et al. 2022). When introduced to Hawaii, it facilitated the transmission of avian malaria and avian poxvirus, which led to the extinction of several endemic bird species, which many more bird species at risk because of the mosquito’s high adaptability and invasive tendencies (Harvey-Samuel et al. 2021). Although there is no evidence that the potential presence of *Cx. quinquefasciatus* in Ireland currently threatens avian fauna, it is wise to monitor its possible emergence and spread. Our findings, together with existing models, highlight the importance of being vigilant about the impacts this invasive mosquito species could have on health and biodiversity, based on its effects elsewhere in the world.

The potential detection of hybrid species also poses concerns. *Culex pipiens pallens* is a widely distributed species in Northern China and a primary vector of lymphatic filariasis and epidemic encephalitis, along with other arboviruses including WNV (Fonseca et al. 2009; Cano et al. 2014; Zhang et al. 2021). Following the large-scale application of insecticides in Northern China, *Cx. pipiens pallens* has also demonstrated a level of insecticide resistance, further hampering control efforts of the species. While this species has not been reported in Europe to date, the potential influence of trading practices between Asia and Europe do pose some concern. For example, Zang et al. (2023) pointed out that the Hunagdao port in China acts as an important node for trade between over 450 ports, including those within Europe. Additionally, it has been proposed that hybrids of *Cx. pipiens* form *pipiens* and *Cx. pipiens* form *molestus* exist and that their combined feeding strategies (i.e. mammalophilic and ornithophilic) could result in species with increased vector transmission capabilities (Huang et al. 2009).

The knowledge of the *Culex pipiens* complex in Ireland and Britain is incomplete. In Ireland, only *Cx. pipiens* was previously recorded (Ashe et al. 1991), whereas in Britain four species have been identified, including *Cx. modestus*, *Cx. territans*, and *Cx. torrentium*. Morphologically distinguishing these species, especially as larval stages, is problematic. For example, some of the larvae initially classified as *Cx. territans* on the basis of an extremely high siphonal index (following Cranston et al. (1987) and Snow (2015)) were, in this study, molecularly identified as *Cx. pipiens* form *pipiens* and *Cx. quinquefasciatus*, suggesting a degree of morphological variation within the *Cx. pipiens* complex at different developmental stages.

Behavioural, physiological, and host preference differences exist among complex members (Zittra et al. 2016), yet their morphological similarity poses identification challenges. Adult differentiation typically relies on the presence of pre-alar scales (frontal wing scales), or wing vein measurement in adults (Börstler et al. 2014; Leggewie et al. 2016), and *Cx. quinquefasciatus* identification often involves examining male genitalia (Harbach 1988; Turell 2012). However, hybridization between *Cx. quinquefasciatus* and *Cx. pipiens* complicates matters further (Sanogo et al. 2008; Farajollahi et al. 2011; Mathews et al. 2017).

### Implications

Proper identification is essential for evaluating mosquito-borne disease risks, given the differing vector capacities within the complex. *Culex pipiens* and *Cx. quinquefasciatus* are known moderate vectors of Sindbis virus (SINV), while *Cx. torrentium* has demonstrated greater susceptibility to SINV and is thought to potentially have a more significant role in the transmission of SINV transmission (Hubálek 2008; Turell 2012; Hesson et al. 2015). Despite humans typically being dead-end hosts, incidences of SINV cases have been acknowledged in Northern Europe since the 1960s, with symptoms like fever and arthralgia (Ling et al. 2019), warrant further investigation of this virus in Ireland.

The coexistence of known and newly reported mosquito species in Ireland raises additional concerns for disease transmission. The presence of both *Cx. torrentium* and *Ae. cinereus* (noted by Ashe et al. 1991) could facilitate a SINV outbreak, with *Cx. torrentium* serving as the primary enzootic host and *Ae. cinereus* as a bridge vector between infected birds and humans (Lundström et al. 1990; Schäfer et al. 2004). Key to such an outbreak are SINV- susceptible birds like *Turdus* spp., *Phylloscopus trochilus*, *Sylvia borin*, and *Erithacus rubecula*, all found in Ireland (Lundström 1999; Schäfer et al. 2004; Kurkela et al. 2008; Bird Watch Ireland 2023).

Additionally, the unique vector capabilities of *Cx. pipiens* ecotypes, which are competent vectors for SINV, WNV, and Usutu virus (USUV), must be considered (Lundström et al. 1990; Vogels et al. 2017; Romo et al. 2018; Camp et al. 2019; Jansen et al. 2019). The ornithophilic nature of *Cx. pipiens* form *pipiens*, predominantly feeding on birds (Becker et al. 2010; Gomes et al. 2013), plays a significant role in WNV transmission, especially in regions where it shows a preference for human hosts, as demonstrated in Central and Mid-Atlantic United States (Andreadis 2012). The detection of this ecotype, alongside *Cx. torrentium*, underscores the potential for increasingly competent vector species in Ireland.

## Conclusions

This is the first molecular examination of Irish mosquito specimens and has revealed a more diverse population than what had previously been reported by Ashe et al. (1991). Based on morphological characterisation, Ashe and colleagues found just one *Culex* species in Ireland, namely *Cx. pipiens*. However, molecular methods employed in the present study facilitated the identification of *Cx. torrentium* and *Cx. pipiens* form *pipiens*. Identifications of these two species was not unexpected and appear reliable. Some of our analysis also revealed the potential presence of *Cx. quinquefasciatus*, but we have some concerns regarding the reference sequences from GenBank used to identify this species.

Based on the results presented in this study, COI is a generally suitable marker for species-level identification while ITS1 can be used to discriminate members of the *Cx. pipiens* complex, including ecotype species. Using the ITS1/2 primers the identification of *Cx. pipiens* form *pipiens* could be further corroborated, but the locus also revealed some ambiguity, particularly in relation to *Cx. quinquefasciatus*. Although this focused on samples collected in the North-East of Ireland and at undisclosed potential points of entry for exotic mosquito species, future studies aiming to reveal mosquito populations, particularly in regions where complexes are known to exist, should employ an island-wide approach to surveillance using a complementary approach involving morphological characterisation and multi-locus sequencing using both mitochondrial and nuclear regions to ensure a comprehensive estimation of mosquito species/sub-species. It is also crucial that suitable and reliable reference libraries are developed for the identification of mosquitoes in Europe, and particularly Ireland, which incorporate local genetic diversity and haplotypes.

### Experimental Procedures Sample Collection

Sampling was carried out in Warrenpoint, Co. Down in Northern Ireland and Dundalk, Co. Louth in the Republic of Ireland (Fig 3). Larvae and adults were collected from plant pots and artificial ponds in an urban location in Warrenpoint (n = 43 larvae (NI labelled samples)) and Dundalk (n = 24 larvae (DL labelled samples); n = 9 adults (DA labelled samples)). Samples were collected using Pasteur pipettes and transferred to a plastic container containing some of the water the larvae were found in. These samples were taken to the laboratory at South East Technological University (SETU) within two days of collection. In the laboratory, larvae or adults were placed individually into a 1 mL tube containing absolute ethanol, to preserve the specimen prior to DNA extraction. In addition, larvae collected by environmental health officers from the HSE at potential points of entry at airports and seaports, at undisclosed locations, and morphologically identified as *Cx. territans* at the National Museum of Ireland (NMI), based on characters in Snow (2015) and Cranston et al. (1987), (n = 8 larvae (NMI labelled samples)), were taken to SETU in absolute ethanol for molecular confirmation of the species.

### DNA Extraction, Polymerase Chain Reaction (PCR), and DNA sequencing

For adult mosquitoes (n = 9), two legs per specimen were used for DNA extraction. This minimally invasive approach was taken to preserve the bulk of the specimen for any future morphological assessments. For larval specimens, those freshly collected and those provided via the NMI (n = 75), DNA was extracted using the whole sample.

DNA was extracted using the NucleoSpin DNA Extraction Kit (Macherey-Nagel) according to manufacturer’s instructions and DNA was eluted in 50 µL of molecular grade water. The concentration (ng/µL) and quality (260/280 ratio) of each DNA extract was measured using a Thermo Scientific NanoDrop™ 8000 Spectrophotometer.

Initially, two genetic regions, COI and ITS1, were used for species identification via PCR and Sanger sequencing. Additionally, when specimens were suspected of belonging to the *Culex pipiens* complex after initial sequencing with COI and ITS1, a third primer set was used to target a fragment of the ITS1/2 locus. This primer set was designed specifically to differentiate species within the *Culex pipiens* complex (Table 2).

**Table 2:**
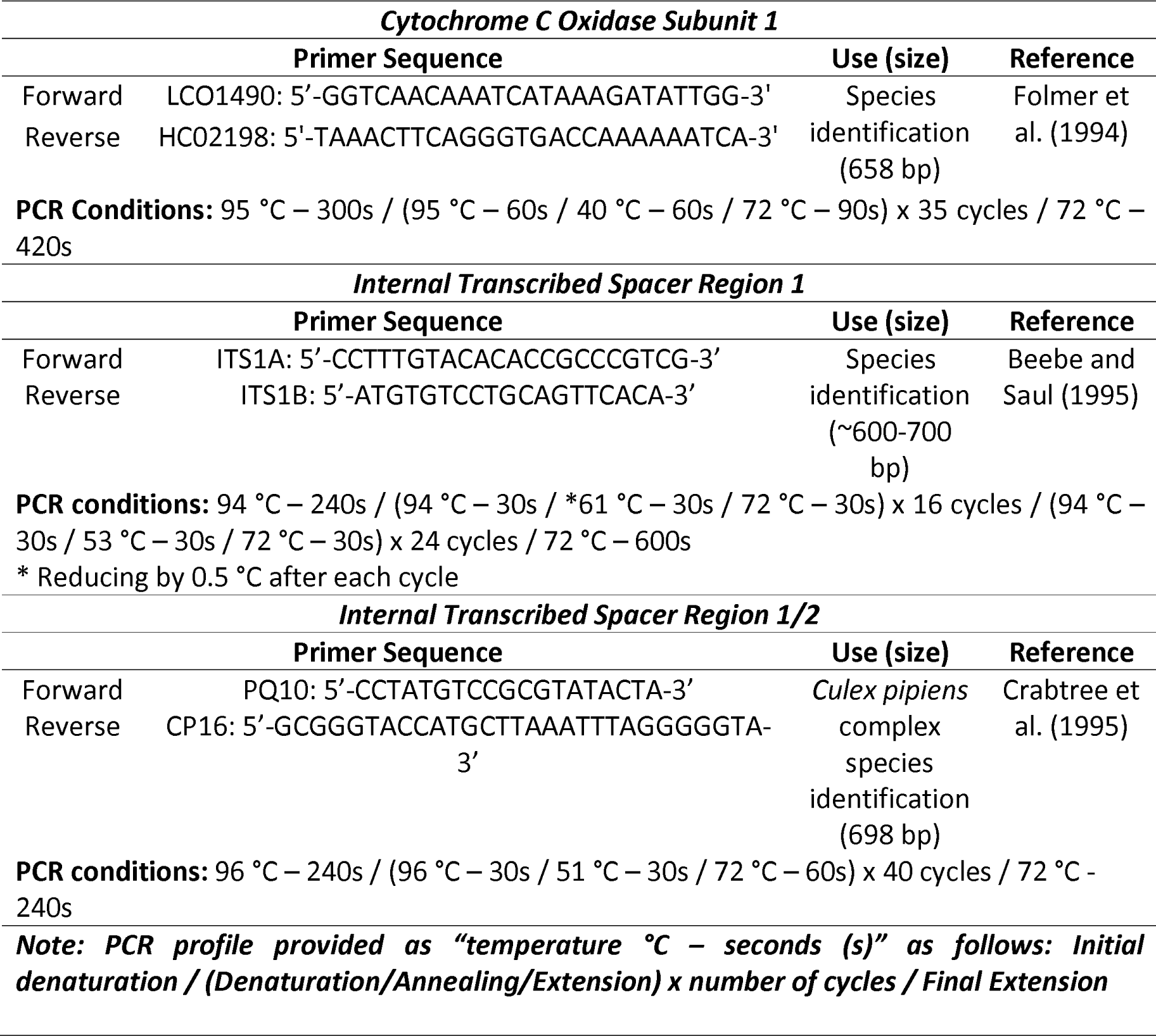
Details of the primer sets used for species and complex identification.

For all protocols, PCR reactions contained 5 µL GoTaq Green HotStart Mastermix, 3 µL molecular grade water, 1 µL template DNA (negative controls contained 1 µL molecular grade water), and 1 µL of the primer mixture containing both the forward and reverse primers (each at a concentration of 50 µM). PCR amplicons were visualised by agarose gel electrophoresis on a 1.6% (w/v) agarose gel.

PCR products were cleaned using MicoCLEAN (Clent Life Science) as per manufacturer’s instructions and sequenced in the forward direction on the Applied Biosystems 3500 Genetic Analyzer using the BigDye Terminator Cycle Sequencing Kit V3.1 (Applied Biosystems) and the standard sequencing POP-7 protocol.

### DNA Sequence Analysis

All sequences were initially assessed for quality in Chromas (Version: 2.6.6) and aligned using ClustalW 1.6 in Molecular Evolutionary Genetics Analysis (MEGA) (Version: 10 (X)) (Kumar et al. 2008). Sequences were then compared to the GenBank repository of nucleotide sequences, and the top hit was recorded along with the sequence accession number, percentage identity, and percentage query cover.

### Analysis of Culex species

The COI and ITS1 *Culex* sequences generated in this study that were identified with a minimum of 98% sequence identity to a GenBank reference species, were included in downstream analyses. However, only sequences representing each of the identified haplotypes were included in the analysis, ensuring the removal of duplicate sequences. Sequences were aligned with the following reference sequences available on GenBank for the species: *Cx. torrentium* (COI accession numbers: HF562558, HF562561, HF562576, HE997145, HE997146 JQ253807, JQ253820, JQ253828, FN395194; ITS1 accession numbers: U33037, U33038, U33039, U33040), *Cx. quinquefasciatus* (COI accession numbers: OP002045, KC250017, MK713993, MN733801, MT108614, MT919714, MT999163; ITS1 accession numbers: U22121, U22122 U22123, U22124, U22125, U22126, U22127, U22128, U22129, U22130, U22131, U22132, MG546064, MG546066, MG546067, JF825532, EU359697, GU562872), *Cx. pipiens* (COI accession numbers: MK971974, MK972000, MK402763, MK402879, MT519668; ITS1 accession number: KU056508), *Cx. pipiens* form *pipiens* (COI accession numbers: JQ253845, JQ253846, JQ253847, KP728854, KP728873, KP728875, KP728876, AM403476; ITS1 accession numbers: U22111, U22112, U22113, U22114, U22117, U22118, U22119, U22120), *Cx. pipiens* form *molestus* (COI accession numbers: KU495006, KU495007, KU495008, KU756486, KJ012149, KJ012153, KJ012156, KP728871, AM403492; ITS1 accession numbers: U22115, U22116), *Cx. pipiens pallens* (COI accession numbers: LC646366, LC646367, LC646368, LC054477, MF179187, KC407754; ITS1 accession numbers: U33025, U33026), and *Cx. territans* (COI accession numbers: KM258162, KM258163, KM258164, KM258165, MW535815, HQ982279, JF873250; ITS1 accession numbers: U33035, U33036) using MEGA. The COI sequences were trimmed to 506 bp. ITS1 sequences trimmed at a matching location resulted in fragments ranging from 320 bp to 355 bp. This variable length is due to the presence of indel (nucleotide insertion/deletions) regions along this locus. Maximum-Likelihood phylogenetic trees were constructed in MEGA using 1000 bootstrap replications and visualised using the Interactive Tree of Life (iTOL: https://itol.embl.de/). For ITS1/2 *Culex* sequences, a Maximum-Likelihood phylogenetic tree was also constructed, based on a 289 bp fragment, using the study samples, the top GenBank identity, and other possible identities suggested by GenBank that had a minimum 99% sequence identity to the sample sequence, and an outgroup representative (the biting midge [Diptera: Certaopogonidae] *Culicoides impunctatus*), was included in that analysis.

