## Supplementary Information for "DNA barcoding reveals an increased diversity within the genus *Culex* (Diptera: Culicidae) in Ireland"

**Supplementary Table 1: A summary of the distribution across the British Isles, habitat preferences and vector capacity of mosquitoes present in the British Isles (WNV: West Nile virus; TAHV: Tahyna virus; SSH: Snowshoe hare encephalitis; INKV: Inkoo virus; DENV: Dengue; CHIKV: Chikungunya fever; RVFV: Rift Valley fever; SINV: Sindbis virus; EEE: Eastern equine encephalitis; LEDV: Lednice virus; USUV: Usutu virus).**

| ***Species*** | ***Distribution*** | ***Preferred habitats*** | ***Vector capacity*** | ***References*** |
| --- | --- | --- | --- | --- |
| *Anopheles algeriensis* | Ireland and Great Britain; rare | Warmer temperatures (thermophilic species) | *Plasmodium* | Medlock et al. (2007); Tippelt et al. (2018) |
| *Anopheles atroparvus** | Great Britain; widespread | Coastal waters, saline and freshwater | *Plasmodium*, WNV, *Dirofilaria* | Medlock et al. (2007); Danabalan et al. (2014); Becker et al. (2010); Filipe (1972); Ferreira et al. (2015); Sainz-Elipe et al. (2010) |
| *Anopheles claviger* | Ireland and Great Britain; widespread | Permanent waters | *Plasmodium* | Medlock et al. (2007); Tippelt et al. (2018); Tagliapietra et al. (2019) |
| *Anopheles daciae** | Great Britain; local reports/suggested to be widespread | Permanent waters | *Plasmodium* (Little is known of the vector capabilities of this species) | Danabalan et al. (2014); Medlock et al. (2007); Rydzanicz et al. (2017); Linton et al. (2005); Culverwell et al. (2020); Brugman et al. (2017); |
| *Anopheles maculipennis* complex* | Ireland and Great Britain; variable (widespread/ local) | Permanent waters | *Plasmodium* | Medlock et al. (2007); White (1978); Linton et al. (2007); Kampen et al. (2016); Kavran et al. (2018) |
| *Anopheles messeae** | Great Britain; widespread | Permanent water, freshwater, brackish waters | *Plasmodium* | Medlock et al. (2007); Bates (1941); Becker et al. (2010); Novikov and Vaulin (2014); Takken et al. (2002); Lindsay et al. (2010) |
| *Anopheles plumbeus* | Ireland and Great Britain; widespread | Tree holes (dendrolimnic), artificial habitats | *Plasmodium*, *Dirofilaria* | Medlock et al. (2007); Schaffner et al. (2012); Becker et al. (2010); Heym et al. (2017); Schaffner et al. (2001) |
| *Aedes annulipes* | Great Britain; widespread | Temporary pools, forests with moist soil | TAHV, Myxoma virus | Medlock et al. (2007); Medlock and Vaux (2015); Foster and Walker (2019); Schäfer and Lundström (2001); Kemenesi et al. (2015); Versteirt et al. (2012) |
| *Aedes cantans* | Ireland and Great Britain; widespread | Woodland pools | WNV, TAHV | Medlock et al. (2007); Service (1977); Service (1973); Szentpali-Gavaller et al. (2014) |
| *Aedes caspius* | Ireland and Great Britain; limited distribution | Coastal waters, inland and coastal marshes, irrigation canals | - | Medlock et al. (2007); Carrieri et al. (2008) |
| *Aedes cinereus* | Ireland and Great Britain; widespread | Flooded habitats | WNV | Medlock et al. (2007); Medlock and Vaux (2009); Andreadis et al. (2004); Timmermann and Becker (2010) |
| *Aedes communis* | Great Britain; uncommon | Temporary pools | SSH, INKV, TAHV | Medlock et al. (2007); Becker and Ludwig (1983); Schäfer and Lundström (2001); Söderström and Nilsson (1987); Belloncik et al. (1982); Lwande et al. (2017); Wegner (2009) |
| *Aedes detritus* | Ireland and Great Britain; widespread | Coastal waters, saline and freshwater | WNV | Medlock et al. (2007); Blagrove et al. (2016); Kampen et al. (2015); Medlock and Vaux (2013); Medlock and Vaux (2015) |
| *Aedes dorsalis* | Ireland and Great Britain; common | Freshwater and brackish water | WNV | Medlock et al. (2007); Rydzanicz et al. (2011); Goddard et al. (2002) |
| *Aedes flavescens* | Ireland and Great Britain; common | Temporary pools, flooded habitats | *Dirofilaria*, TAHV*,* equine arboviruses | Medlock et al. (2007); Horsfall et al. (1958); Söderström and Nilsson (1987); Ibanez-Justicia et al. (2015); Medlock and Vaux (2014); Versteirt et al. (2012); Chapman et al. (2016); Frimeth and Arai (1983); Hubalek and Halouzka (1997); Ibanez-Justicia et al. (2015) |
| *Aedes geminus* | Great Britain; unsure but likely common | Flooded habitats | WNV | Medlock et al. (2007); Medlock and Vaux (2009); Andreadis et al. (2004); Timmermann and Becker (2010) |
| *Aedes geniculatus* | Great Britain; common | Tree holes (dendrolimnic) | *Dirofilaria*, WNV | Medlock et al. (2007); Bockova et al. (2013); Mughini-Gras et al. (2013) |
| *Aedes luecomelas* | Great Britain; few reports | Wet/flooded meadows and swamps | WNV, DENC, CHIKV, *Plasmodium* | Medlock et al. (2007); Rydzanicz et al. (2011); Schäfer et al. (2006); Ibanez-Justicia et al. (2015); Medlock and Vaux (2015); Stroo et al. (2018) |
| *Aedes nigrinus* | Great Britain; common | Small ponds; flooded meadows | No vector capabilities have been identified | Medlock et al. (2007); Harbach et al. (2017); Ibanez-Justicia et al. (2015) |
| *Aedes punctor* | Ireland and Great Britain; common | Woodland pools | *Wolbachia* | Medlock et al. (2007); Ricci et al. (2002); Medlock et al. (2005) |
| *Aedes rusticus* | Great Britain; common | Flooded habitats, temporary groundwater | RVFV | Medlock et al. (2007); Lumley et al. (2018); Rey et al. (2001) |
| *Aedes sticticus* | Great Britain; historical reports | Temporary pools, wetlands | *Dirofilaria*, RVFV | Medlock et al. (2007); Schäfer et al. (2008); Buxton and Mullen (1980); Iranpour et al. (2011) |
| *Aedes vexans* | Great Britain; common | Temporary pools, rivers, marshes | WNV, RVFV, SINV, EEE | Medlock et al. (2007); Joe and Clay (2002); Medlock et al. (2017); Cupp et al. (2004); Gendernalik et al. (2017); Goddard at al. (2002); Lilja et al. (2018); Talla et al. (2016) |
| *Coquillettidia richiardii* | Ireland and Great Britain; widespread | Permanent waters | WNV | Medlock et al. (2007); Schäfer and Lundström (2001); Sérandour et al. (2006); Komar (2000) |
| *Culex modestus* | Great Britain; historical reports | Marshes, pond edges, rice fields | WNV, LEDV, TAHV | Medlock et al. (2007); Medlock and Vaux (2012); Pradel et al. (2009); Bakonyi et al. (2013); Berčič et al. (2019); Ponçon et al. (2007); Danielova and Holubova (1977) |
| *Culex pipiens^^^* | Ireland and Great Britain; widespread | Underground and permanent surface waters | USUV, WNV, RFVF, SINV, TAHV, *Dirofilaria* | Medlock et al. (2007); Dehghan et al. (2014); Osório et al. (2014); Martinet et al. (2019) |
| *Culex territans* | Great Britain; common | Swamps; marshes; springs; wetland pools | WNV, EEE, frog erythrocytic virus | Medlock et al. (2007); Kitron and Pener (1986); Joe and Clay (2002); Bartlett-Healy et al. (2008); Burkett-Cadena et al. (2008); Gruia-Gray and Desser (1992) |
| *Culex torrentium^^^* | Great Britain; common | Permanent water, flooded habitats, artificial habitats | SINV, WNV | Medlock et al. (2007); Dehghan et al. (2010); Scherpner (1960); Weitzel et al. (2015); Jöst et al. (2010); Leggewie et al. (2016) |
| *Culiseta alaskaensis* | Ireland and Great Britain; common | Open pools and semi-permanent waters | Has not been linked to any diseases | Medlock et al. (2007); Ashe et al. (1991); Kampen et al. (2013); Schäfer and Lundström (2001) |
| *Culiseta annulata* | Ireland and Great Britain; widespread | Still water; artificial environments | WNV, RVFV | Medlock et al. (2007); Hamidian (2013); Brugman et al. (2017); Reeves et al. (2016); Toma et al. (2008) |
| *Culiseta fumipennis* | Great Britain; widespread | Permanent waters, temporary pools within woodlands | No vector capabilities have been identified for this species | Medlock et al. (2007); Medlock and Vaux (2015); Schäfer and Lundström (2001); Snow and Medlock (2008); Chapman et al. (2016); Medlock et al. (2005) |
| *Culiseta litorea* | Ireland and Great Britain; widespread | Coastal water | WNV | Medlock et al. (2007); Schäfer and Lundström (2001); Medlock and Vaux (2015). Service (1994) |
| *Culiseta longiareolata* | Great Britain; common | Artificial habitats; pools and ditches | WNV. Some studies have suggested no vector capability of this species. | Medlock et al. (2007); Becker et al. (2010); Seidel et al. (2013); Hubalek and Halouzka (1999); Romi et al. (2004); Engler et al. (2013); Rioz et al. (2007) |
| *Culiseta morsitans* | Ireland and Great Britain; widespread | Permanent waters and forests | WNV | Medlock et al. (2007); Schäfer and Lundström (2001); Medlock and Vaux (2015). Service (1994) |
| *Culiseta subochrea* | Ireland and Great Britain; uncommon | Permanent waters, flooded habitats, artificial habitats | Little is known about the vector capabilities of this species | Medlock et al. (2007); Snow and Medlock (2008); Shaalan et al. (2017) |
| *Orthopodomyia pulcripalpis* | Great Britain; uncommon | Tree holes (dendrolimnic) | Assumed vector of avian arboviruses | Medlock et al. (2007); Medlock et al. (2005); Becker et al. (2010); Zittra et al. (2017) |
| **** These species form the Anopheles maculipennis complex***  ***^ These species form the Culex pipiens complex***  ***_ Species known to occur in Ireland***  ***~ Information largely from UK-based studies as only one Irish study focused on mosquitoes has been performed ~*** | | | | |

***Supplementary Table 2: Average DNA concentrations and standard deviations for mosquito adults and larvae.***

| **Sample set** | **N** | **Average DNA concentration (ng/µL)** | **Standard Deviation (**±) |
| --- | --- | --- | --- |
| Dundalk Larvae (DL) | 24 | 123.84 | 74.66 |
| Dundalk Adults (DA) | 9 | 8.28 | 4.41 |
| Northern Ireland Larvae (NI) | 43 | 93.36 | 65.26 |
| National Museum of Ireland Larvae (NMI) | 8 | 2.2 | 1.31 |
| **Material Type** | **N** | **Average DNA concentration (ng/µL)** | **Standard Deviation (**±) |
| Larvae | 75 | 93.39 | 73.16 |
| Adults | 9 | 8.28 | 4.41 |
| **All samples** | **84** | **84.27** | **73.99** |

**Supplementary Table 3: Species identification of samples collected from Northern Ireland (NI), Dundalk (DA = Dundalk Adults; DL = Dundalk Larvae), and from the Health Service Executive but provided by the National Museum of Ireland (NMI) using the optimised sequencing methodologies for the COI and ITS1 gene regions. Sequences highlighted represent Culex species sequences that had a minimum of 98% sequence identity and were used for downstream analysis.**

|  | ***Cytochrome C Oxidase Subunit 1*** | | | ***Internal Transcribed Spacer Region 1*** | | |
| --- | --- | --- | --- | --- | --- | --- |
| ***Sample*** | ***%ID/Query*** | ***Identification*** | ***bp*** | ***%ID/Query*** | ***Identification*** | ***bp*** |
| ***NI 1*** | 99.84/99 | *Cx. torrentium* (MH463062) | 633 | 88.46/62 | *Cx. torrentium* (U33039) | 579 |
| ***NI 2*** | 99.68/97 | *An. claviger* (KM457607) | 650 | 100/ 36 | *An. algeriensis* (MT808464) | 369 |
| ***NI 3*** | 99.69/99 | *Cx. torrentium* (MH463062) | 651 | 84.59/97 | *Cx. torrentium* (U33039) | 585 |
| ***NI 4*** | 100/96 | *An. claviger* (MK403247) | 656 | 99.26/36 | *An. algeriensis* (MT808464) | 364 |
| ***NI 5*** | 99.84/96 | *An. claviger* (OK465168) | 647 | 99.26/36 | *An. algeriensis* (MT808464) | 365 |
| ***NI 6*** | 99.69/99 | *Cx. torrentium* (MH463062) | 649 | 82.55/98 | *Cx. torrentium* (U33039) | 557 |
| ***NI 7*** | 100/100 | *Cx. pipiens* (MH463070) | 649 | 87.50/100 | *Cx. pipiens pipiens* (U22117) | 542 |
| ***NI 8*** | 100/96 | *An. claviger* (MK403247) | 647 | 99.26/36 | *An. algeriensis* (MT808464) | 364 |
| ***NI 9*** | 99.24/100 | *Cx. torrentium* (FN395799) | 658 | 98.50/98 | *Cx. torrentium* (U33039) | 676 |
| ***NI 10*** | 99.13/100 | *An. claviger* (MK403538) | 575 | 99.26/36 | *An. algeriensis* (MT808464) | 367 |
| ***NI 11*** | 87.48/97 | *Tavastia sp.* (HQ105367 | 649 | 97.12/32 | *Clunio tsushimensis* (AB704963) | 548 |
| ***NI 12*** | 99.38/100 | *Cx. torrentium* (FN395194) | 646 | 93.32/98 | *Cx. torrentium* (U33039) | 611 |
| ***NI 13*** | 99.84/98 | *Cx. torrentium* (MH463062) | 645 | 93.31/98 | *Cx. torrentium* (U33039) | 645 |
| ***NI 14*** | 100/99 | *Cx. torrentium* (MH463062) | 640 | 98.92/98 | *Cx. torrentium* (U33038) | 656 |
| ***NI 15*** | 100/99 | *Cx. torrentium* (MH463062) | 648 | 93.00/93 | *Cx. torrentium* (U33039) | 361 |
| ***NI 16*** | 99.84/99 | *Cx. torrentium* (MH463062) | 638 | 79.97/98 | *Cx. torrentium* (U33039) | 580 |
| ***NI 17*** | 99.84/98 | *Cx. torrentium* (MH463062) | 648 | 81.44/86 | *Cx. torrentium* (U33039) | 581 |
| ***NI 18*** | 99.87/99 | *Cx. torrentium* (MH463062) | 641 | 94.20/97 | *Cx. torrentium* (U33039) | 497 |
| ***NI 19*** | 100/99 | *Cx. torrentium* (MH463062) | 638 | 94.72/98 | *Cx. torrentium* (U33039) | 533 |
| ***NI 20*** | 99.84/99 | *Cx. torrentium* (MH463062) | 638 | 93.77/45 | *Cx. torrentium* (U33039) | 570 |
| ***NI 21*** | 100/99 | *Cx. torrentium* (MH463062) | 649 | 90.00/99 | *Cx. torrentium* (U33039) | 569 |
| ***NI 22*** | 99.84/99 | *Cx. torrentium* (MH463062) | 645 | 98.52/99 | *Cx. torrentium* (U33039) | 679 |
| ***NI 23*** | 99.39/99 | *Cx. torrentium* (FN395194) | 659 | 93.98/99 | *Cx. torrentium* (U33039) | 468 |
| ***NI 24*** | 99.85/99 | *Cx. torrentium* (MH463062) | 649 | 82.10/97 | *Cx. torrentium* (U33039) | 526 |
| ***NI 25*** | 99.84/98 | *Cx. torrentium* (MH463062) | 645 | 93.49/45 | *Cx. torrentium* (U33039) | 570 |
| ***NI 26*** | 99.84/99 | *Cx. torrentium* (MH463062) | 637 | 93.41/100 | *Cx. torrentium* (U33039) | 637 |
| ***NI 27*** | 99.85/99 | *Cx. torrentium* (MH463062) | 649 | 94.29/100 | *Cx. torrentium* (U33039) | 578 |
| ***NI 28*** | 99.85/100 | *Cx. torrentium* (MH463062) | 647 | 93.56/99 | *Cx. torrentium* (U33039) | 640 |
| ***NI 29*** | 99.85/100 | *Cx. torrentium* (MH463062) | 646 | 82.27/86 | *Cx. torrentium* (U33039) | 579 |
| ***NI 30*** | 100/98 | *Cx. torrentium* (MH463062) | 646 | 86.49/100 | *Cx. torrentium* (U33038) | 442 |
| ***NI 31*** | 99.69/100 | *Cx. torrentium* (MH463062) | 644 | 89.53/65 | *Cx. torrentium* (U33039) | 552 |
| ***NI 32*** | 100/97 | *An. claviger* (MK403247) | 644 | 100/ 35 | *An. algeriensis* (MT808464) | 380 |
| ***NI 33*** | 99.69/99 | *Cx. torrentium* (MH463062) | 649 | 85.50/99 | *Cx. torrentium* (U33039) | 612 |
| ***NI 34*** | 100/97 | *An. claviger* (MK403247) | 643 | 98.52/35 | *An. algeriensis* (MT808464) | 378 |
| ***NI 35*** | 100/97 | *An. claviger* (MK403247) | 644 | 100/ 35 | *An. algeriensis* (MT808464) | 379 |
| ***NI 36*** | 99.69/99 | *Cx. torrentium* (MH463062) | 641 | 83.45/97 | *Cx. torrentium* (U33039) | 569 |
| ***NI 37*** | 100/97 | *An. claviger* (MK403247) | 647 | 99.26/35 | *An. algeriensis* (MT808464) | 379 |
| ***NI 38*** | 100/97 | *An. claviger* (MK403247) | 648 | 100/ 33 | *An. algeriensis* (MT808464) | 365 |
| ***NI 39*** | 100/97 | *An. claviger* (MK403247) | 644 | 100/ 35 | *An. algeriensis* (MT808464) | 380 |
| ***NI 40*** | 100/97 | *An. claviger* (MK403247) | 637 | 99.26/36 | *An. algeriensis* (MT808464) | 364 |
| ***NI 41*** | 99.84/97 | *An. claviger* (OK465168) | 643 | 99.26/36 | *An. algeriensis* (MT808464) | 368 |
| ***NI 42*** | Sample not included | | | Sample not included | | |
| ***NI 43*** | 100/97 | *An. claviger* (MK403247) | 644 | 99.26/35 | *An. algeriensis* (MT808464) | 379 |
| ***DA 1*** | 100/99 | *Cx. torrentium* (MH463062) | 641 | 85.22/100 | *Cx. torrentium* (U33038) | 607 |
| ***DA 2*** | 99.42/100 | *Cs. annulata* (MT192948) | 521 | 99.26/34 | *Ae. mcintoshi* (KJ940824) | 621 |
| ***DA 3*** | 99.83/99 | *Cs. annulata* (LC476724) | 598 | 98.50/33 | *Cs. novaezealandiae* (MG546077) | 615 |
| ***DA 4*** | 100/100 | *Cx. pipiens* (MK714013) | 628 | 99.45/99 | *Cx. quinquefasciatus* (EU359697) | 723 |
| ***DA 5*** | 100/99 | *Cs. annulata* (LC476724) | 644 | 99.21/33 | *Ae. metallicus* (KU056497) | 612 |
| ***DA 6*** | 100/98 | *Cx. pipiens* (MK714013) | 645 | 98.75/100 | *Cx. pipiens pipiens* (U22113) | 641 |
| ***DA 7*** | 100/99 | *Cx. pipiens* (MK714012) | 644 | 80.07/98 | *Cx. pipiens pipiens* (U22113) | 602 |
| ***DA 8*** | 100/99 | *Cx. torrentium* (MH463062) | 622 | 94.06/99 | *Cx. torrentium* (U33038) | 439 |
| ***DA 9*** | 99.68/100 | *Cx. pipiens* (MN460845) | 618 | 92.74/100 | *Cx. pipiens pipiens* (U22113) | 565 |
| ***DL 1*** | 99.38/99 | *Cs. annulata* (LC476717) | 651 | 99.26/25 | *Ae. mcintoshi* (KJ940824) | 542 |
| ***DL 2*** | 100/100 | *Cx. pipiens* (MK714012) | 639 | 96.38/99 | *Cx. pipiens pipiens* (U22117) | 581 |
| ***DL 3*** | 99.69/99 | *Cs. annulata* (LC476724) | 650 | 98.50/32 | *Cs. novaezealandiae* (MG546077) | 624 |
| ***DL 4*** | 98.76/99 | *Chydorus sphaericus* (MT872704) | 649 | Poor sequence quality | | |
| ***DL 5*** | 100/99 | *Cs. annulata* (LC476727) | 649 | 98.50/33 | *Cs. novaezealandiae* (MG546077) | 615 |
| ***DL 6*** | 99.84/100 | *Cx. pipiens* (MH463059) | 641 | 85.39/99 | *Cx. pipiens pipiens* (U22113) | 571 |
| ***DL 7*** | 100/99 | *Cx. pipiens* (MH463059) | 642 | 95.30/99 | *Cx. pipiens pallens* (U33026) | 686 |
| ***DL 8*** | 99.84/100 | *Cx. pipiens* (MK714012) | 630 | 87.82/89 | *Cx. quinquefasciatus* (MG546066) | 649 |
| ***DL 9*** | 99.84/98 | *Cx. pipiens* (MH463059) | 649 | 96.21/99 | *Cx. pipiens* (KU056508) | 750 |
| ***DL 10*** | 99.84/98 | *Cx. torrentium* (MH463062) | 645 | 97.41/98 | *Cx. torrentium* (U33040) | 701 |
| ***DL 11*** | 100/98 | *Cx. pipiens* (MH463070) | 648 | 97.93/100 | *Cx. quinquefasciatus* (MG546066) | 627 |
| ***DL 12*** | 99.85/100 | *Cx. pipiens* (MH463070) | 649 | 94.30/98 | *Cx. pipiens pipiens* (U22113) | 603 |
| ***DL 13*** | 100/98 | *Cx. pipiens* (MK714012) | 646 | 92.27/90 | *Cx. pipiens pipiens* (U22113) | 600 |
| ***DL 14*** | 100/100 | *Cx. pipiens* (MN460845) | 558 | 93.60/99 | *Cx. pipiens pipiens* (U22113) | 659 |
| ***DL 15*** | 100/99 | *Cx. pipiens* (MH463070) | 641 | 97.34/99 | *Cx. pipiens pipiens* (U22113) | 637 |
| ***DL 16*** | 99.85/100 | *Cx. pipiens* (MH463070) | 649 | 95.67/64 | *Cx. pipiens pipiens* (U22117) | 501 |
| ***DL 17*** | Poor sequence quality | | | Poor sequence quality | | |
| ***DL 18*** | 100/98 | *Cx. pipiens* (MH463070) | 648 | 99.87/99 | *Cx. pipiens pipiens* (U22113) | 748 |
| ***DL 19*** | 99.85/98 | *Cx. pipiens* (MH463059) | 657 | 93.01/100 | *Cx. pipiens pipiens* (U22113) | 601 |
| ***DL 20*** | 9100/100 | *Cx. pipiens* (MH463070) | 632 | 94.44/100 | *Cx. pipiens pallens* (U33025) | 615 |
| ***DL 21*** | 100/100 | *Cx. pipiens* (MH463070) | 641 | 97.84/100 | *Cx. quinquefasciatus* (EU359697) | 681 |
| ***DL 22*** | 100/98 | *Cx. pipiens* (MK714013) | 645 | 99.33/100 | *Cx. pipiens pipiens* (U22114) | 744 |
| ***DL 23*** | 100/100 | *Cx. pipiens* (MH463070) | 640 | 98.76/100 | *Cx. pipiens pipiens* (U22117) | 679 |
| ***DL 24*** | 100/99 | *Cx. pipiens* (MH463070) | 641 | 99.36/100 | *Cx. quinquefasciatus* (MG546066) | 627 |
| ***NMI 1*** | 99.37/97 | *Cx. pipiens* (KJ401308) | 648 | 95.28/100 | *Cx. pipiens pipiens* (U22113) | 439 |
| ***NMI 2*** | 100/100 | *Cx. pipiens* (MH463070) | 643 | 92.98/96 | *Cx. pipiens pipiens* (U22114) | 370 |
| ***NMI 3*** | 99.84/98 | *Cx. pipiens* (KJ401308) | 640 | 94.39/98 | *Cx. pipiens pipiens* (U22113) | 518 |
| ***NMI 4*** | 100/100 | *Cx. quinquefasciatus* (MK370091) | 387 | 93.00/88 | *Cx. pipiens pipiens* (U22114) | 400 |
| ***NMI 5*** | 100/100 | *Cx. pipiens* (HG793473) | 561 | 87.38/86 | *Cx. pipiens pipiens* (U22114) | 375 |
| ***NMI 6*** | 95.48/99 | *Cx. pipiens* (KJ401308) | 623 | 100/40 | *Cx. quinquefasciatus* (MG546067) | 359 |
| ***NMI 7*** | 100/97 | *Cx. pipiens* (KJ401308) | 594 | 92.31/81 | *Cx. pipiens pipiens* (U22114) | 400 |
| ***NMI 8*** | 99.32/100 | *Cx. pipiens* (MT199095) | 589 | 97.36/99 | *Cx. pipiens* (KU056508) | 749 |
|  | *Indicate samples with poor sequence quality* | | | | | |

**Supplementary Table 4: Sequence IDs of COI sequences that were listed as being identical via the PopART software (names highlighted in red refer to samples from this study).**

| ***Node ID*** | ***Identical Sequences*** |
| --- | --- |
| NMI3 | NMI5 NMI7 |
| CxPipPip_KP728876 | CxPipPip_KP728873 CxPipPip_KP728875 CxPipPip_KP728854 CxPipMol_KJ012149 CxPipMol_KJ012161 CxPipMol_AM403492 CxPipMol_KJ012153 CxPipPal_LC646368 CxPipPal_MF179187 CxPipPal_KC407754 |
| NI7 | DA4 DA6 DA7 DA9 DL2 DL7 DL8 DL11 DL12 DL13 DL14 DL15 DL16 DL18 DL19 DL20 DL21 DL22 DL23 DL24 NMI2 CxPip_MH463070 CxPip_MK714013 CxPip_MK714012 CxPip_MN460845 CxPip_MK603829 CxPip_MK403381 CxPip_MH463059 CxPip_MK714001 CxPip_HG793489 CxPip_MK713992 CxPip_MK713986 CxPip_HG793437 CxPip_MK713985 CxPip_MK713995 CxPip_MK713988 CxPip_KX260940 CxPip_HE997152 CxPip_MK971888 CxPip_MK972009 CxPip_MT519630 CxPip_MK402879 CxPip_MT519668 CxPip_MK402763 CxPip_MK971974 CxPip_MK972000 CxQuin_MK713993 CxQuin_KC250017 CxQuin_OP002045 CxQuin_MN733801 CxQuin_MT919714 CxQuin_MT108614 CxPipPip_AM403476 CxPipPip_JQ253847 CxPipPip_JQ253846 CxPipPip_JQ253845 CxPipMol_KU495007 CxPipMol_KU756486 CxPipMol_KU495008 CxPipMol_KU495006 CxPipPal_OK493372 CxPipPal_LC646367 CxPipPal_LC646366 CxPipPal_LC054477 |
| NI1 | NI14 NI15 NI19 NI21 NI30 DA1 DA8 CxTorr_MH463062 CxTorr_FN395199 CxTorr_HE997135 CxTorr_MK403031 CxTorr_MK971897 CxTorr_HE997076 CxTorr_HE997144 CxTorr_MK971841 CxTorr_MK971836 CxTorr_HE997137 CxTorr_MK971795 CxTorr_HF562558 CxTorr_HE997145 CxTorr_JQ253820 |
| NI3 | NI6 NI9 NI31 NI33 NI6 |
| NI12 | NI13 NI16 NI17 NI18 NI20 NI22 NI23 NI24 NI25 NI26 NI27 NI28 NI29 DL10 CxTorr_FN395194 |
| CxTerr_KM258164 | CxTerr_MW535815 |

**Supplementary Table 5: Sequence IDs of ITS1 sequences that were listed as being identical via the PopART software (names highlighted in red refer to samples from this study).**

| **Node ID** | **Identical Sequences** |
| --- | --- |
| CxQuin_EU359697 | CxQuin_MG546067 CxPip_KU056508 |
| DA4 | CxQuin_U22124 CxPipPip_U22120 |
| DA6 | DL18 DL22 DL23 DL24 CxQuin_U22130 CxQuin_U22129 CxPipPip_U22113 CxPipPip_U22114 CxPipPal_U33026 CxPipPal_U33025 |
| CxPipPip_U22117 | CxPipPip_U22118 |
| CxQuin_U22132 | CxQuin_U22122 |
| CxPipPip_U22112 | CxPipPip_U22111 |
| NI9 | NI14 NI22 |
| CxPipMol_U22115 | CxPipMol_U22116 |
| CxPipXCxQuin_U33044 | CxPipXCxQuin_U33043 |
